# Identification of antineoplastic agents for oral squamous cell carcinoma: an integrated bioinformatics approach using differential gene expression and network biology

**DOI:** 10.1101/2021.10.14.464376

**Authors:** Abdussalam Usman, Faisal F. Khan

## Abstract

Oral squamous cell carcinoma (OSCC) is the most common malignant epithelial neoplasm and anatomical subtype of head and neck squamous cell carcinoma (HNSCC) with an average 5-year survival rate of less than 50%. To improve the survival rate of OSCC, the discovery of novel anti-cancer drugs is urgently needed. In the present study, we performed metanalysis of 5 gene expression datasets (GSE23558, GSE25099, GSE30784, GSE37991 and TCGA-OSCC) that resulted in 1851 statistically significant DEGs in OSCC. The DEGs were involved in key biological pathways that drive the progression of OSCC. A comprehensive protein-protein interaction (PPI) network was constructed from the DEGs and the top protein clusters (modules) were extracted in Cytoscape. The DEGs from the top modules were searched for antineoplastic agents using L1000CDS^2^ server. The search resulted in a total of 37 perturbing agents from which 12 well-characterized antineoplastic agents were selected. The selected 12 antineoplastic agents namely Teniposide, Palbociclib, Etoposide, Fedratinib, Tivozanib, Afatinib, Vemurafenib, Mitoxantrone, Idamycin, Canertinib, Dovitinib and Selumetinib. These drugs showed interactions with the over expressed hub genes that regulate cellular proliferation and growth in OSCC progression. These identified antineoplastic agents are candidates for their potential role in treating OSCC.

## Introduction

Oral Squamous Cell Carcinoma (OSCC) is the most common oral malignancy which accounts for approximately 90% of all malignant neoplasms of the oral cavity [1, 2]. OSCC is a subtype of Head and Neck Squamous Cell Carcinoma (HNSCC) that arises in the squamous lining of several sites of oral cavity i.e. tongue, lip, gingiva, palate, floor of the mouth and buccal mucosa [3, 4]. OSCC is the 16^th^ most common malignancy around the globe with nearly 1 million prevalent cases in 2020 [5]. The risk factors that are increasing the prevalence of OSCC around the globe are tobacco smoking and chewing, betel quid, alcohol, poor oral hygiene and periodontitis [6]. The current available treatment modality of OSCC are surgical resection, radiotherapy, chemotherapy and targeted therapy however, the 5-year survival rate of approximately 50% is still unchanged [7–10].

The survival rate of OSCC patients can be improved through the development of novel drugs with the help of next-generation technologies and distinguished status of a tumor [11, 12]. However, the translation of the novel drugs into clinical practice have several challenges i.e. lower success rate in clinical trials, high cost and time scales that can span more than a decade [13, 14].To overcome some of these challenges, drug repositioning or repurposing uses an alternative strategy for identification of new uses i.e. the existing drugs that are outside scope of the original medical indication [15, 16]. The key advantage of drug repositioning is that the pharmacodynamic, pharmacokinetic and toxicity profiles of drugs have already been established in the original preclinical and Phase-I studies. These drugs could therefore be rapidly progressed into Phase-II and Phase-III clinical studies and the associated development cost and time could be significantly reduced [17, 18]. Drug repositioning has been made feasible with publicly accessible databases and repositories that store information of gene expression and biological pathways in cancer [19, 20] in human cell lines or model organisms exposed to perturbing agents or small molecules [21–23]. The application of biomarkers is not only restricted to therapeutic targeting but can also be used for diagnosis or prognosis of a cancer patient [24, 25]. The library of integrated network-based cellular signatures (LINCS) is one such project that generates response gene signatures induced by a diverse collection of perturbing agents in different model systems such as cell lines, differentiated cells and embryonic stem cells [23, 26]. The correlation of expression signatures induced by drugs or perturbing agents with the gene expression profile of a cancer offers an efficient strategy for repurposing existing drugs for new indications [27, 28]. The correlation of drugs and cancer can be established by deploying several computational methods such as systems biology, bioinformatics, machine learning and network biology for drug repositioning in complex diseases such as cancer and in identification of new indications existing drugs [29].

The computational approach for drug repositioning mostly utilizes a “guilt by association” strategy of correlating small molecules or existing drugs with the disease due to regulation of similar biological pathways and gene expression [30]. These correlations between disease and drugs can be predicted based on similar chemical structures, genetic variations and gene expression profiles [31]. Currently, the interest in the use of transcriptomics-guided drug repositioning is expanding and so far, a number of studies have been conducted [32]. Transcriptomics-guided drug repositioning in cancer is more feasible because of limited background knowledge of cancer type and drugs is require [33, 34].

Recently, transcriptomics-guided drug repositioning by targeting the overly expressed genes and enriched pathways have been conducted in different types of cancer such as breast cancer [35, 36], prostate cancer [37, 38], gastric cancer [39], HNSCC [40] and many others. However, there is still no such study available that explored drug repositioning for OSCC by using gene expression data. In the present study, we performed differential gene expression analysis of multiple OSCC datasets and identified a set of genes that were driving tumorigenesis and the progression of OSCC. The gene clusters were inversely correlated with the antineoplastic signatures retrieved from the L1000CDS^2^ server.

## Materials and Methods

### Searching for gene expression datasets

A comprehensive search for gene expression datasets was performed in Gene Expression Omnibus (GEO) – an online public repository hosted by NCBI for genomics and transcriptomics datasets (http://www.ncbi.nlm.nih.gov/geo/). The key searching terms for identification of datasets of interest in GEO database were “*Oral squamous cell carcinoma*”, “*OSCC*”, “*Oral cancer*” and “*Oral cavity cancer*” and the search was conducted in August of 2020. The results were further filtered by selecting terms such as “*Homo sapiens*” as a host and expression profiling by array and “*expression profiling by high throughput sequencing*” as a study type. From the resulting list, only those datasets were selected that resulted from the original experimental studies and also included the data of gene expression from OSCC samples as well as controls. In addition, OSCC samples were also extracted from The Cancer Genome Atlas (TCGA-HNSC) mRNA expression dataset accessed using the Broad GDAC Firehose (https://gdac.broadinstitute.org/).

### Preprocessing of the datasets

The selected datasets were characterized by retrieving information such as GEO accession, GEO platform (GPL) ID, number of OSCC cases and controls, sample type and gene expression data were extracted. The differential expression analysis of datasets retrieved from the GEO and TCGA were performed by using the limma and DESeq2 package in R v3.6 respectively. From the individual datasets, the significant differentially expressed genes (DEGs) were detected by adjusting the p-value to correct the false positive results by using Benjamini & Hochberg false positive discovery rate method [41]. The volcano plot for each dataset was created by using “EnhancedVolcano” package in R. The DEGs were filtered on the basis of adjusted p-value (adj. *P*) < 0.01 and the probe set IDs in different datasets were reannotated into gene symbols by using the “biomartr” package in R. The number DEGs detected in each dataset were visualized as a Venn Diagram using InteractiVenn tool [42].

The mean log_2_ fold change (log_2_FC) was estimated where more than one probe set IDs were representing the same gene symbol. By merging the DEGs from all datasets, the log_2_FC of the DEGs were estimated by taking the mean value of the DEGs that were already filtered on the basis of adj. *P* < 0.05.

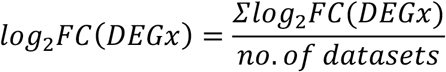

Where log_2_FC of each DEG was estimated by taking mean of log_2_FC that was found in datasets after applying cutoff of adj. *P* > 0.05. The DEGs were further filtered based on mean log_2_FC > |1| and detection in at least 3 datasets (Supplementary Table 1).

### Gene ontology and KEGG terms enrichment analysis

The annotation, visualization and integrated discovery (DAVID) database (https://david.ncifcrf.gov/) was employed for enrichment analysis of gene ontology (GO) and Kyoto Encyclopedia of Genes and Genomes (KEGG) terms. The results from the submitted listed of DEGs were filtered using a cutoff of p-value < 0.05 and the background reference was set to the whole genome annotation for human genome. The significantly enriched top ten GO and KEGG terms were plotted as barplot by using the “ggplot2” package in R.

### Construction of PPI and the detection of modules

The protein-protein interaction (PPI) network was built by submitting the list of DEGs in Search Tool for the Retrieval of Interacting Genes (STRING v11; “https://string-db.org/”). The PPI network was constructed by setting a cutoff criterion for confidence score of > 0.4 for the nodes and the data was imported into Cytoscape v3.7.1 for visualization. The network file was imported into Cytoscape and network components which had less than 5 nodes were removed. For further exploration of densely interacting modules in the network, the Molecular Complex Detection (MCODE) plugin in Cytoscape which detects modules or densely interacting nodes in the network. The default parameters such as degree cutoff = 2, node score cutoff = 0.2 and k-Core of 2 in MCODE were applied. For identification of enriched pathways regulated by the DEGs in top modules, the ClueGO app of Cytoscape was employed for identification of enriched KEGG pathways with P-value < 0.05 as a threshold.

### Correlation between DEGs of top modules and drugs

Drug repositioning was performed by targeting the DEGs in top 5 modules via L1000CDS^2^. L1000CDS^2^ is an online search tool that has knowledge about upregulated and downregulated genes perturbed in the cell lines after treated with drug or perturbagens. L1000CDS^2^ utilizes L1000 data from the Library of Integrated Network-based Cellular Signatures (LINCS) and return results of top 50 perturbations for the submitted query. L1000CDS^2^ was employed by submitting the genes in modules that showed mean log2FC > 1 as upregulated and log2FC < - 1 as downregulated genes (Supplementary Table 2). The parameters were set to default and enabled the search for expressional signatures of small molecules that were in negative correlation (reversed) to the input. The submitted query resulted into a total of 50 chemical perturbations and their corresponding heatmap was constructed by using Pheatmap package in R.

### Screening for antineoplastic agents

The resulting list of L1000CDS^2^ was screened for antineoplastic agents by retrieving information i.e. drug classification, mechanism of action (MoA), existing indication, targeting pathways and clinical trials from PubChem (https://pubchem.ncbi.nlm.nih.gov/). Only those antineoplastic agents were considered significant which are currently under investigation in phase-III of clinical trials or approved by FDA.

### Construction of drug-gene interaction network

The identified antineoplastic agents were searched for targeting genes by employing Drug Gene Interaction Database (DGIdb) and the list of genes were extracted. A network of drugs targeting genes was built and visualized in Cytoscape. In the drug-gene network, the overlapping genes from DGIdb and L1000CDS^2^ were also identified. Hub genes were identified by using network analysis tool in the Cytoscape and the nodes were filtered based on degree value ≥10. The role of hub genes in OSCC progression was explored by KEGG enrichment analysis using the DAVID server. The differential expression of hub genes was analyzed using the Gepia2 server [43].

## Results

### Five OSCC gene expression datasets with control samples were identified for the meta-analysis

The gene expression datasets of OSCC were searched in GEO repository by using relevant keywords in search bar and the datasets were further filtered by selecting *homo sapiens* as a specie and gene expression studies either from microarray or high throughput technology. The search resulted into a total of 188 datasets which were further screened for studies that included OSCC samples along with healthy control and had sample size >30. From the filtered result list, only four datasets; GSE23558, GSE25099, GSE30784 and GSE37991 fulfilled the above-mentioned criteria. In addition, OSCC samples from TCGA-HNSC mRNA datasets were also extracted on the basis of anatomical sites of oral cavity that included oral tongue, floor of mouth, buccal mucosa, hard palate and alveolar ridge sites. The characteristics of the selected datasets have been summarized in Table 1 with a total of 575 samples (443 OSCC and 132 controls).

**Table 1.** The selected gene expression datasets.

| Datasets ID | GPL ID | Platform | Sample size |  |
| --- | --- | --- | --- | --- |
|  |  |  | OSCC | Control |
| GSE30784 | GPL570 | Affymetrix Human Genome U133 Plus 2.0 Array | 167 | 45 |
| GSE23558 | GPL6480 | Agilent-014850 Whole Human Genome Microarray 4x44K | 27 | 5 |
| GSE25099 | GPL5175 | Affymetrix Human Exon 1.0 ST Array | 57 | 22 |
| GSE37991 | GPL6883 | Illumina HumanRef-8 v3.0 expression beadchip | 40 | 40 |
| TCGA-OSCC | NA | Illumina HiSeq | 152 | 20 |

### Preliminary analysis found 1851 DEGs that were significantly dysregulated in OSCC

Significant DEGs were identified in the selected gene expression datasets by correlating OSCC with control samples. The differential expression of probe set ids in each dataset have been visualized as volcano plot in Figure 1. The probe set ids were converted into gene symbols and then filtered by setting a cutoff of adj. *P* <0.01. The screening of DEGs based on adj. *P* <0.05 cutoff led to the detection of 3972, 9942, 11666, 9624 and 1762 DEGs in GSE23558, GSE25099, GSE30784, GSE37991 and TCGA-OSCC datasets respectively. The similarities of the DEGs in selected datasets have been illustrated as a Venn diagram as shown in Figure 2A. The DEGs in the datasets were merged on the basis of similar gene symbols and their mean log_2_FC was calculated. For further analysis, DEGs that showed mean log_2_FC > |1| and were detected in at least 3 datasets were considered (Supplementary Table 1). Finally, a total 1851 DEGs were detected which included 741 were upregulated and 1110 were downregulated DEGs.

**Figure 1.**
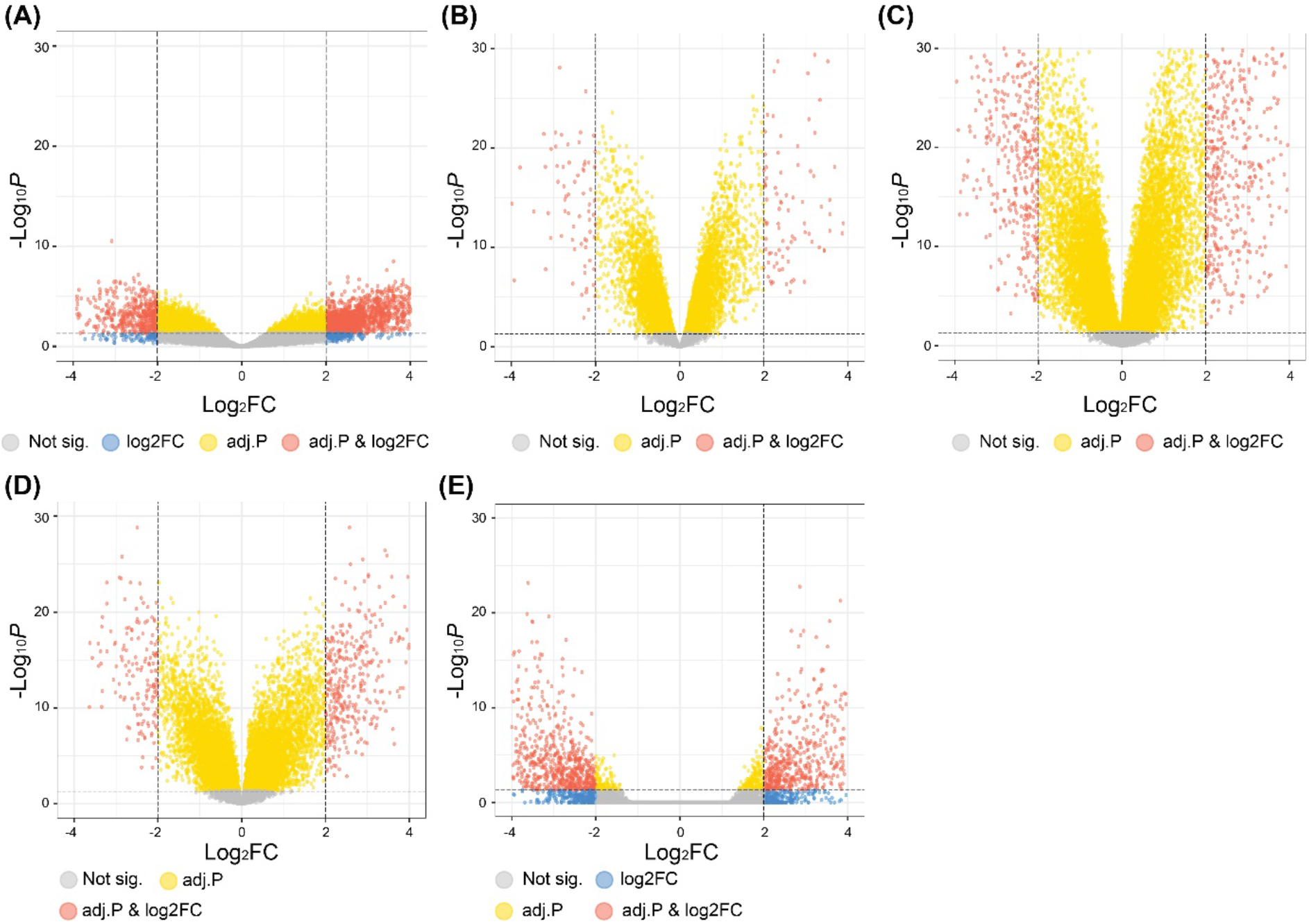
Volcano plots of the selected genes expression dataset using EnhancedVolcano package in R. The significant differentially expressed probes in dataset (A) The differential expression of 3972 probes in GSE23558, (B) 9942 in GSE25099, (C) 11666 in GSE30784, (D) 9624 in GSE37991 and (E) 1762 in TCGA-OSCC after initial processing.

**Figure 2.**
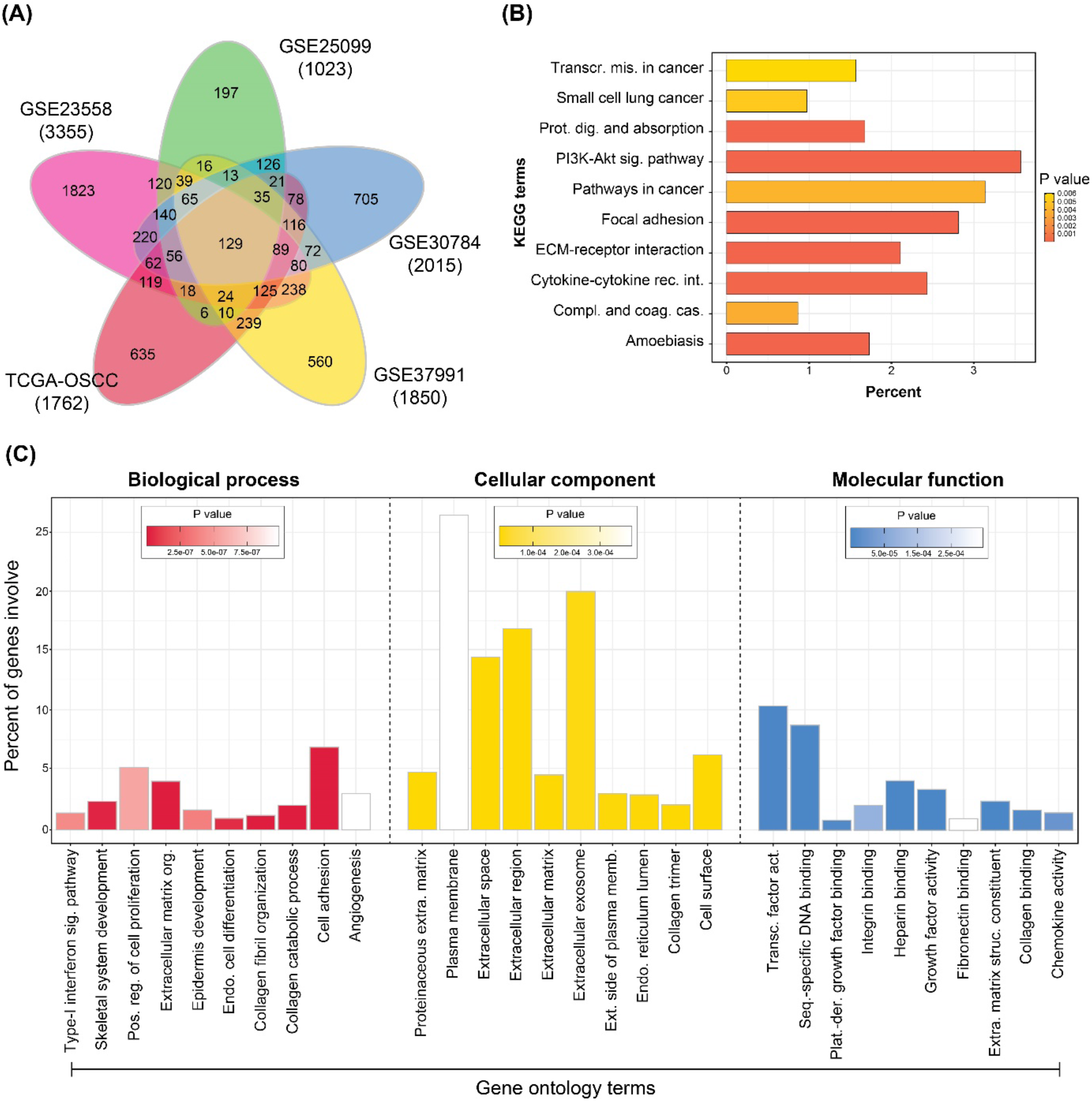
Functional annotation of filtered DEGs from OSCC datasets. (A) Venn diagram of the DEGs identified in each dataset of OSCC using adj. *P* <0.05 and log2FC >|1| as cutoff. (B) Enrichment analysis of KEGG terms in OSCC DEGs identified using the DAVID server. (C) GO terms enrichment analysis of OSCC DEGs using the DAVID server.

### DEGs of OSCC were predominantly involved in cellular proliferation and tumor microenvironment

The enrichment of GO terms in biological process (BP), cellular component (CC) and molecular function (MF) as well as KEGG terms were searched for DEGs by using the DAVID webserver. For identification of statistically significant GO terms, a cutoff of P-value <0.05 was applied which detected a total of 229 BP terms, 64 CC terms and 71 MF terms (Supplementary Table 3). The top ten GO terms for each category have been shown in Figure 1C. The enriched BP terms included the extracellular matrix organization, collagen catabolic process, cell differentiation, cell adhesion, angiogenesis, inflammation response and regulation of cell proliferation. The CC terms included extracellular region, plasma membrane, cell surface and cellular organelles. GO terms for MF included growth factor, monooxygenase and chemokine activity, heparin and metallic ions, collagen and receptor binding. In addition, the top KEGG pathway terms were enriched in PI3K-Akt signaling, cytokine-cytokine receptor interaction, ECM receptor interaction, focal adhesion, transcription misregulation in cancer and pathways in cancer. The top ten enriched KEGG pathways are shown in Figure 1B.

### The PPI network of DEGs had a total of 1777 nodes and 324 nodes that were densely connected

The resulting network from the STRING database showed enrichment p-value <1e-16 and a total of 55 disconnected nodes were removed from the network. The network was visualized in the Cytoscape as shown in Figure 3A. The PPI network showed a total of 1777 nodes and 16958 edges in which 711 nodes were overexpressed and 1048 under-expressed while the expression of 18 nodes was unknown. After MCODE analysis in Cytoscape, the top five densely connected clusters or modules were extracted which are summarized in Table 2.

**Figure 3.**
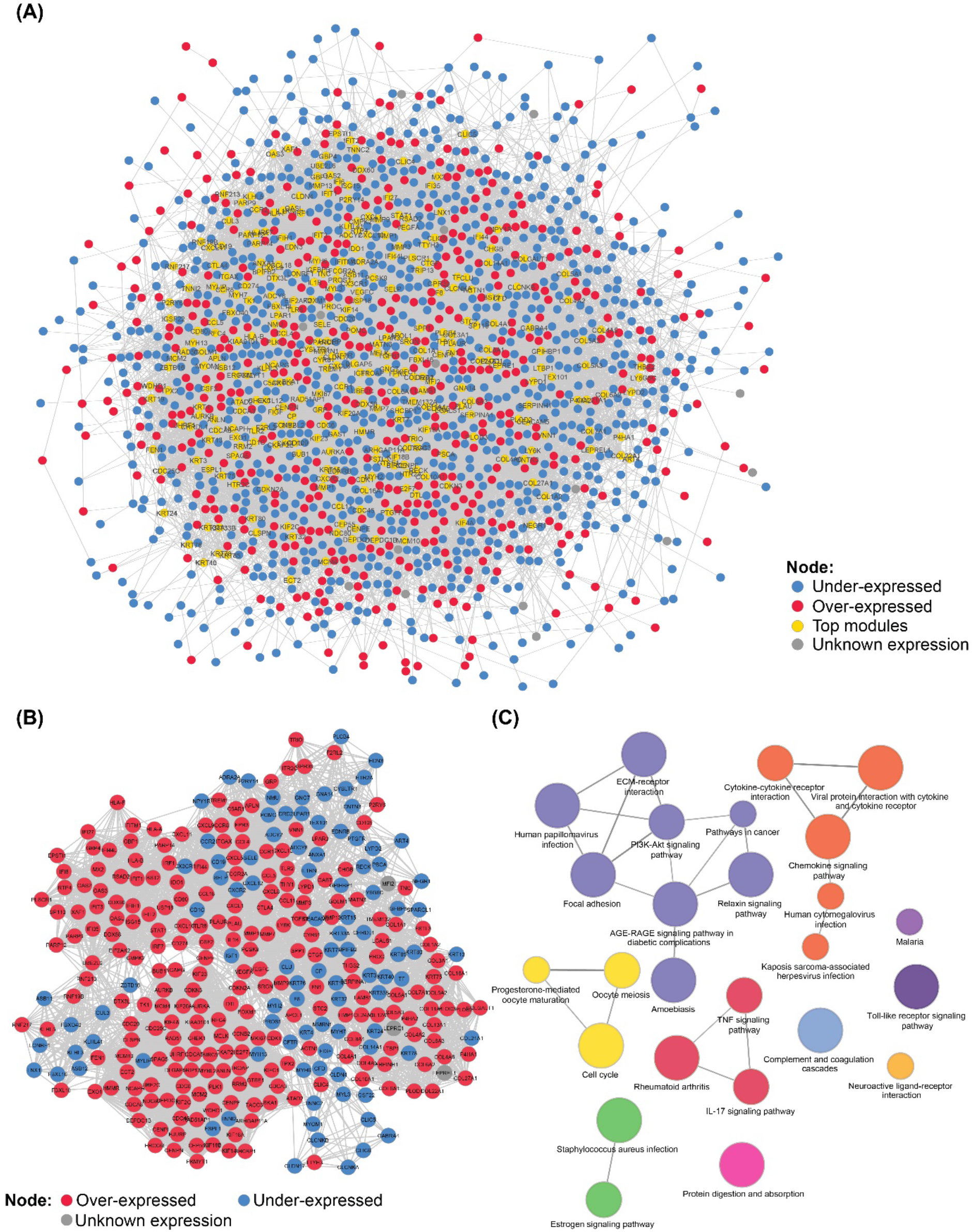
Protein-protein interaction (PPI) network of differentially expressed genes (DEGs) in OSCC. (A) A comprehensive PPI network of DEGs in OSCC. (B) Extracted top 5 modules from the OSCC network using MCODE in Cytoscape. (C) Enrichment of KEGG terms in DEGs from top modules using ClueGO in Cytoscape.

**Table 2.** DEGs summary of each dataset. DEGs from each dataset with cutoff of adj. P < 0.05 and log2FC > |1|

| Dataset | DEGs | uDEGs | dDEGs |
| --- | --- | --- | --- |
| GSE30784 | 2015 | 1028 | 987 |
| GSE23558 | 3355 | 1323 | 2032 |
| GSE25099 | 1023 | 521 | 502 |
| GSE37991 | 1850 | 736 | 1114 |
| TCGA-OSCC | 1762 | 788 | 974 |

The sub-network of the top modules comprised of 323 nodes and 6314 edges in which 224 nodes were overexpressed and 96 under-expressed and 3 were unknown (Supplementary Table 4) and has been shown in Figure 3B. The enrichment analysis of KEGG pathways in the top modules was performed using ClueGO in Cytoscape and the results are as Figure 3C. The terms that were significantly enriched in the top modules include regulating focal adhesion, PI3K-Akt signaling pathway, ECM-receptor interaction, HPV infection, cell cycle, pathways in cancer, TNF signaling pathway, IL-17 signaling pathway, chemokine signaling pathway and cytokine-cytokine receptor interaction.

### Twelve FDA-approved anti-neoplastic agents were found for top DEGs in OSCC

L1000CDS^2^ was used for the identification of candidate drugs by inversely correlating the expression of L1000 signatures in cell lines perturbed by chemical agents against the DEGs in the top modules. The search resulted in identification of top 50 perturbations by 37 candidate drugs. The perturbation of candidate drugs was identified in 11 cell line types i.e. A375, A549, BT20, HCC515, HEPG2, HME1, HT29, LNCAP, MCF7, MCF10A and VCAP. The primary source of these cell lines was breast cancer, lung adenocarcinoma, prostate cancer, hepatoblastoma, malignant melanoma and colon adenocarcinoma. The overlapping score for the perturbation was in the range of 0.19 to 0.23 which is estimated based on the cosine distance between the submitted list of DEGs and L1000 signatures [44]. The expression of L1000 signatures perturbed by the candidate drugs were mostly detected in the A375 and MCF7 cell lines. The dose concentration of the drugs in cell lines were in the range of 0.08 - 80.0 micromolar (uM) and the expression recording time was 24 hours. The expression of L1000 signatures detected in different cell lines that were perturbed by the candidate drugs have been shown as a clustergram in Figure 4.

**Figure 4.**
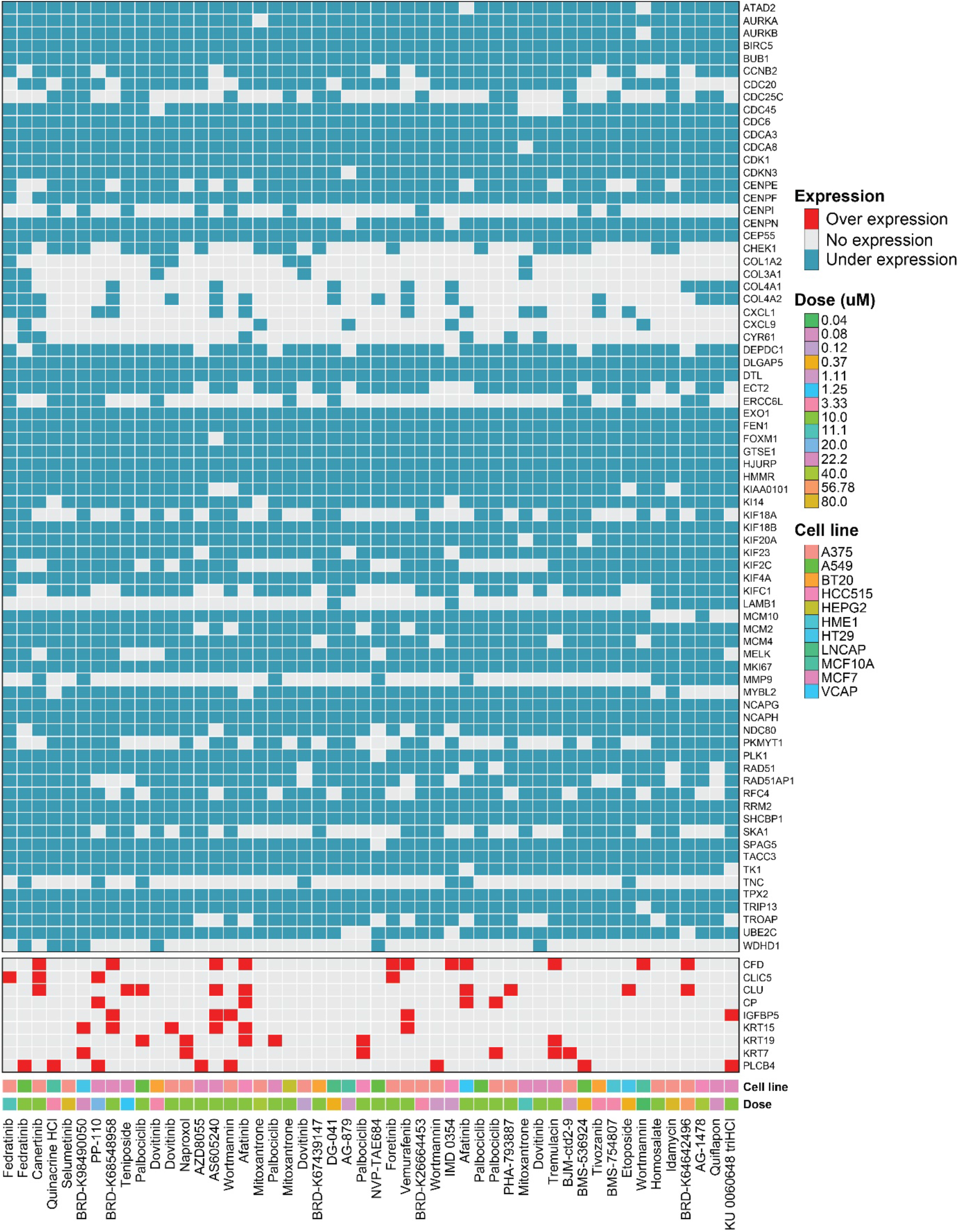
Clustergram of the candidate drugs identified in L1000CDS^2^ database. The reversal expression of the candidate drugs at the bottom row to the input DEGs at the right column have been shown as blue for downregulated and red for upregulated.

The candidate drugs were characterized by retrieving information from the PubChem ChEMBL databases (Supplementary Table 5). Out of 37 drugs, 3 were not found in the PubChem and 7 were poorly characterized in the database and therefore, they were removed from further analysis. The characterized list of 27 drugs had 23 antineoplastic agents and the remaining 4 were classified as anti-inflammatory, anticoagulant and sun screening agents. The 23 antineoplastic agents were further screened for identification of effective antineoplastic agent that are currently in the phase-III of trials or approved by FDA that resulted into 12 drugs and have been characterized in Table 4. Nine antineoplastic agents; Teniposide, Palbociclib, Etoposide, Fedratinib, Tivozanib, Afatinib, Vemurafenib, Mitoxantrone and Idamycin have been approved while three (Canertinib, Dovitinib and Selumetinib) are still in Phase-III of clinical trials.

**Table 3.**
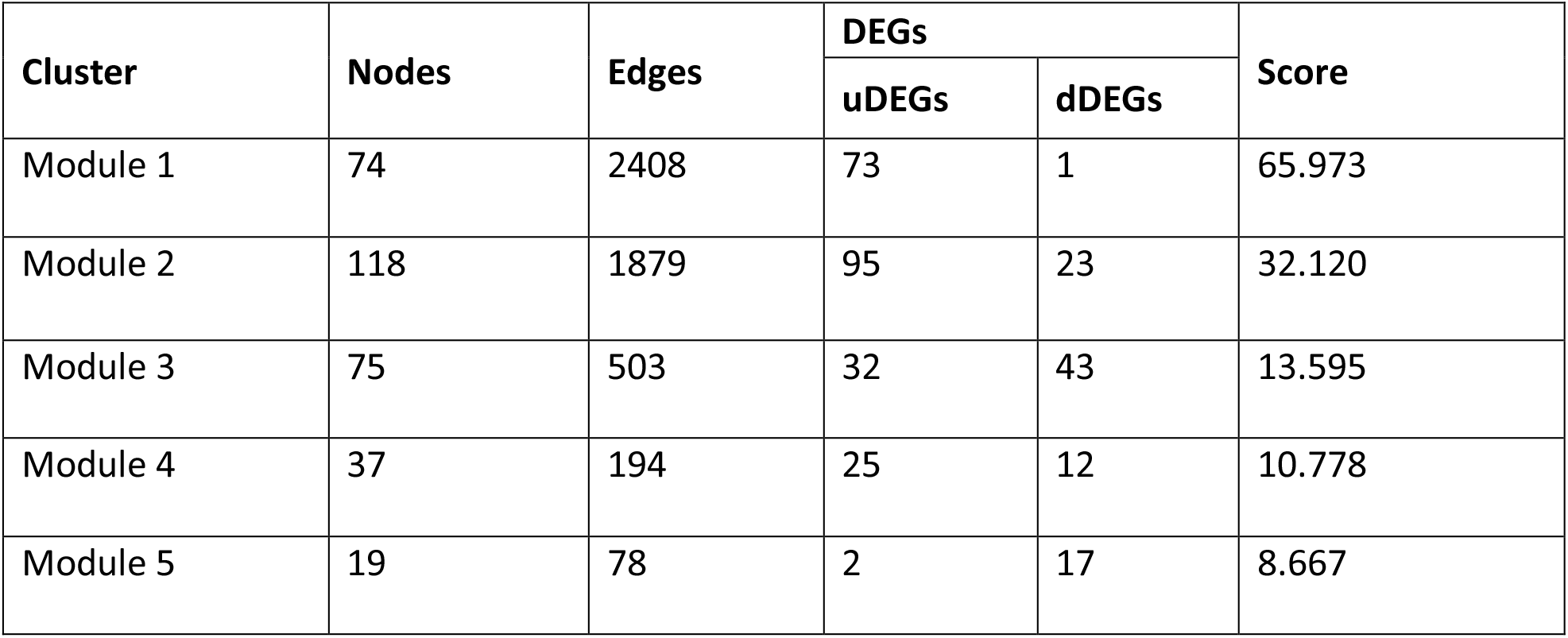
Characteristics of top 5 modules from the PPI network.

| Cluster | Nodes | Edges | DEGs |  | Score |
| --- | --- | --- | --- | --- | --- |
|  |  |  | uDEGs | dDEGs |  |
| Module 1 | 74 | 2408 | 73 | 1 | 65.973 |
| Module 2 | 118 | 1879 | 95 | 23 | 32.120 |
| Module 3 | 75 | 503 | 32 | 43 | 13.595 |
| Module 4 | 37 | 194 | 25 | 12 | 10.778 |
| Module 5 | 19 | 78 | 2 | 17 | 8.667 |

**Table 4.**
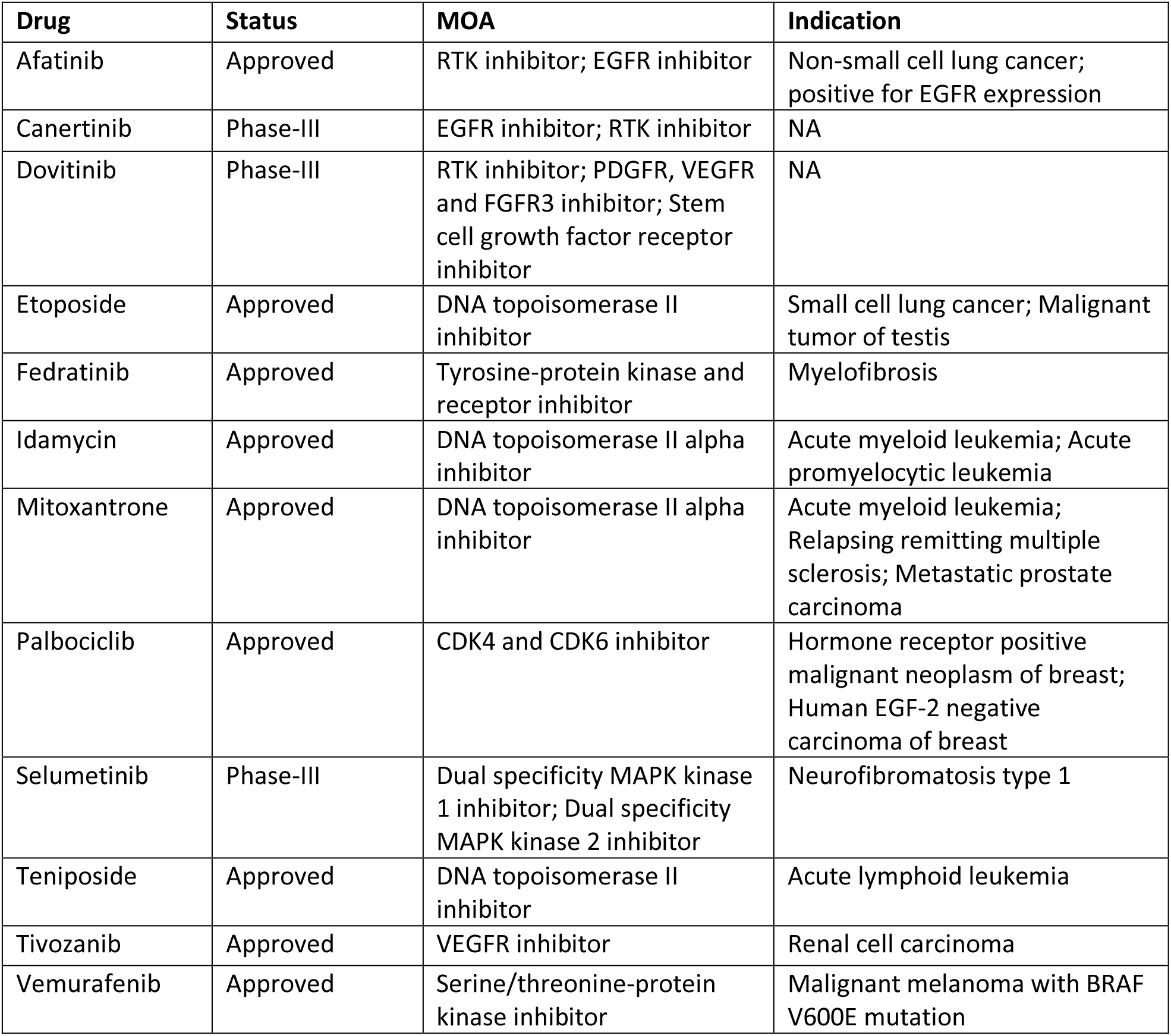
Characteristics of the antineoplastic agents.

The antineoplastic agents were further characterized by exploring their mechanism of action MoA) and indications from PubChem and DrugCentral 2021 database [45]. The MoA of selected candidates were involved in inhibition of cyclin dependent kinase 4 (CDK-4) and 6 (CDK-6), DNA topoisomerase II, mitogen activated protein kinase (MAPK), serine/threonine kinase (STK) and blocking of receptor tyrosine kinases (RTK) such as epidermal growth factor receptor (EGFR), vascular endothelial growth factor receptor (VEGFR), platelet derived growth factor receptor (PDGFR), fibroblast growth factor receptor (FGFR), stem cell growth factor receptor (SCGFR). These antineoplastic agents have been used in the treatment of breast cancer, prostate cancer, renal cell carcinoma, non-small and small cell lung cancer, myelofibrosis, acute myeloid and lymphoid leukemia and neurofibromatosis.

### Construction of a Drug-Gene Network using DGIdb data identified potential targets that regulate cellular proliferation and tumor growth

To further validate, the targeting genes by 12 antineoplastic agents were retrieved from DGIdb and the resulting list was imported and visualized in the Cytoscape. The network displayed antineoplastic agents showing 1119 interactions with 348 genes. The most interacting antineoplastic agents were Dovitinib, Palbociclib and Etoposide that showed interactions with 163, 117 and 112 genes respectively. The nodes of the network were labelled with the expression profile of the antineoplastic agents that resulted earlier from L1000CDS^2^ database as shown in Figure 5A. The integrated network displayed 93 over expressed and 14 under expressed genes in OSCC that were targeted by the antineoplastic agents. The expression of the OSCC genes was annotated from the expression of the genes in the top modules. In addition, the network also included 240 genes that showed interactions with the candidate drugs however, they were not matched with the expression of the genes in the top modules.

**Figure 5.**
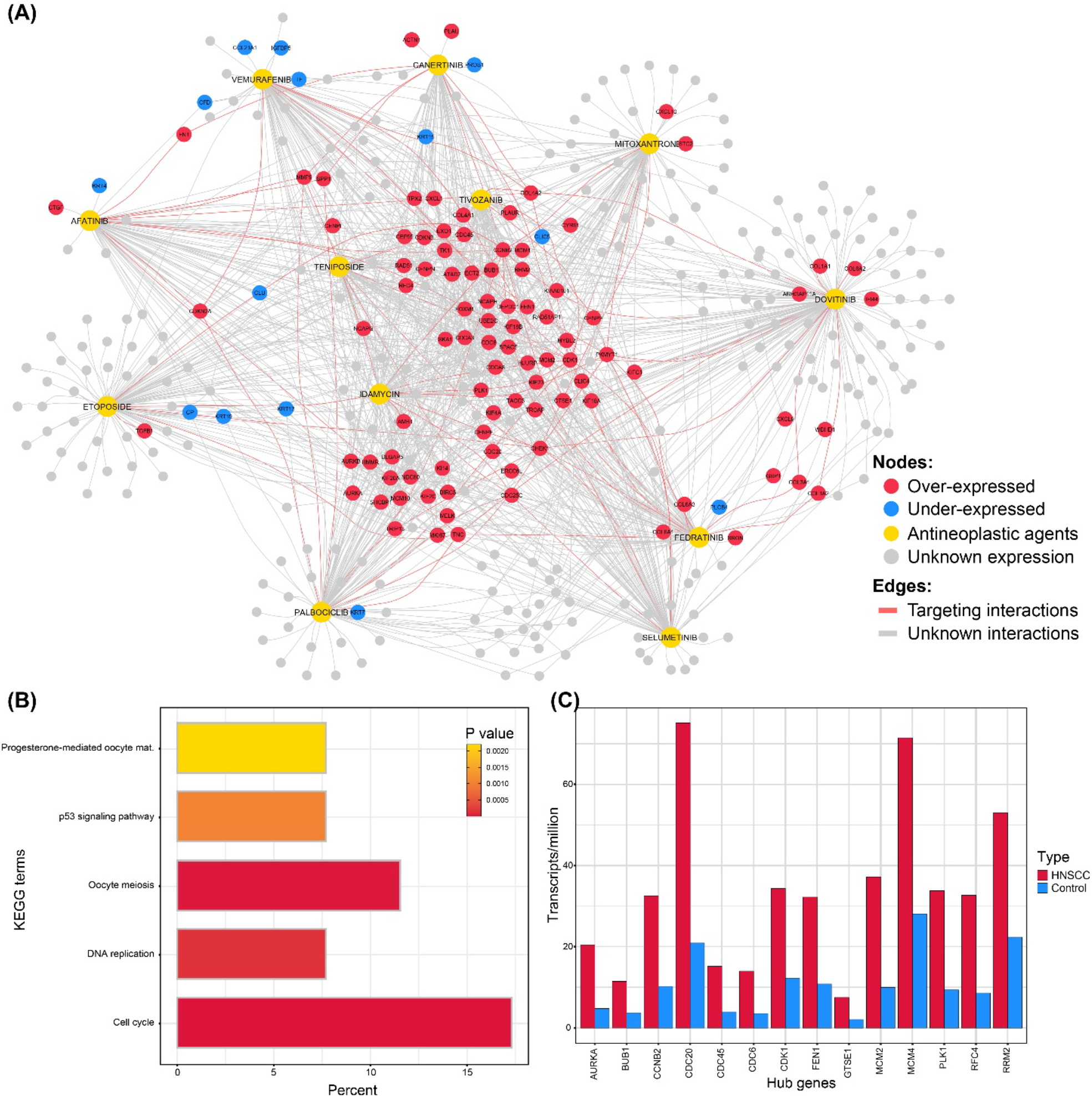
The integrative network of drug gene interaction with expressional signatures from L1000CDS^2^. (A) The 12 well-characterized antineoplastic agents and its targeting interactions with overly-expressed genes in OSCC. (B) The enrichment of the targeting genes and enriched KEGG terms identified from the DAVID server with p-value <0.05 cutoff. (C) The identified core genes from the integrative network and its over-expression in HNSCC datasets using the Gepia2 server.

Hub genes in the drug-gene network were identified by using a cutoff of ≥10 degree in the network which resulted in to a total of 53 genes (Supplementary Table 6). The 53 genes were overexpressed in OSCC and the enrichment analysis showed 14 genes that showed enriched KEGG terms of Cell cycle, DNA replication, p53 signaling pathway, oocyte meiosis and progesterone mediated oocyte maturation as shown in Figure 5B. The 14 hub genes were CCNB2, FEN1, CDC20, RFC4, RRM2, PLK1, CDC45, MCM4, CDK1, MCM2, GTSE1, BUB1, AURKA and CDC6. The overexpression of these genes was verified from Gepia2 server HNSCC mRNA expression dataset from TCGA and genotype-tissue expression (GTEx) as shown in Figure 5C

## Discussion

Drug repositioning is an alternative approach to the traditional drug development offering speed, cost effectiveness, and relative safety for patients going into trials [46]. With the advent of transcriptomics technologies for quantification of gene expression in cancer as well as transcriptional response to small molecular treatment and the availability of the public databases has made the application of drug repositioning more feasible [47, 48]. Currently, the trend in deploying computational or *in-silico* methods for transcriptome-based drug repositioning is increasing and many novel agents have been identified by inverse correlating the gene expression profile of small molecular agents with the cancer [9, 49, 50]. In the present study, we utilized an *in-silico* approach for drug repositioning based on transcriptomic profile of OSCC retrieved from GEO and TCGA repositories.

Several microarray gene expression datasets were found in GEO database that includes samples from OSCC as well as control however, in the case of GSE30784 that also included 17 oral dysplasia samples which were removed from further analysis. For TCGA-HNSC gene expression dataset, the OSCC samples were extracted based on the anatomical site of the tumor. Several packages of R software i.e. MetaDE [51], MetaRE [52] and metaRNASeq [53] are available to perform meta-analysis of gene expression datasets. However due to the differences in the platform of the selected datasets, DEGs were initially filtered from each dataset based on adj. *P* value and then the lists were merged and mean log2FC for each gene was estimated. The DEGs were further filtered by considering mean log2FC >|1| and duplication in ≥3 datasets. The resulting list included a total of 2107 DEGs in which 821 were upregulated and 1286 downregulated.

The enrichment analysis of the filtered DEGs revealed the regulation of pathways associating with cell communication and signaling pathways i.e. focal adhesion, PI3K-Akt signaling pathway, cytokine-cytokine receptor pathways, complement and coagulation cascades, transcriptional misregulation in cancer, pathways in cancer and ECM-receptor interaction. The overexpression of these pathways has been recently explored in OSCC patients [54–57] as well as in HNSCC [40, 58]. These enriched pathways have been linked with local invasion, lymph node metastasis, tumor growth and tumor microenvironment that drives tumorigenesis, cellular proliferation and OSCC progression [59–64].

A PPI network from the list of DEGs was constructed using interaction from the STRING database for identification of highly interacting protein clusters or modules that are involved in molecular pathways necessary for tumor development in OSCC. Within the network, the densely connected or highly interacting protein clusters were identified which resulted into a total of 323 proteins. These proteins were involved in the upregulation of pathways that are involve in tumor microenvironment, inflammatory response, HPV infection, cell proliferation, local invasion and metastasis in OSCC. The enrichment of these pathways in OSCC have been already described in the above paragraph and these pathways drive the main phenotype and progression of OSCC.

For targeting these molecular entities and inhibiting the activity of the key pathways, the list of proteins was selected for drug repositioning by identification of reversal expression form drug expression. The reversal expression of cell lines retrieved from LINCS data in L1000CDS2 which utilizes the cosine distance for inverse correlation between submitted DEGs and stored signatures from drug perturbation. A total of 37 candidate drugs were identified which showed considerable overlap score. The perturbations were observed from 11 types of cell lines and various dose ranges and intervals. The list of drugs was further filtered based on their characterization in PubChem and only antineoplastic agents were selected that resulted in a total of 23 candidates. The selected candidates were screened for 12 well-characterized agents; Teniposide, Palbociclib, Etoposide, Fedratinib, Tivozanib, Afatinib, Vemurafenib, Mitoxantrone, Idamycin, Canertinib, Dovitinib and Selumetinib that were either approved by FDA or in phase-III of clinical trials. The MoA of the selected antineoplastic agents were inhibition of CDK proteins, DNA topoisomerase II, MAPK, STK, EGFR, VEGFR, FGFR and SCGFR which showed similarity with the biological pathways regulated by the DEGs in OSCC.

Eight out of 12 selected anti-neoplastic agents have been already tested for efficiency and toxicity in treating small group of OSCC or HNSCC patients and xenograft models. Teniposide [65] and Vemurafenib [66] have shown potential results in treating small group of OSCC patients. Afatinib [67] and Etoposide [68] also revealed significant out comes of the advanced or metastatic HNSCC patient treatment while Palbociclib [69], Dovitinib [70], Canertinib [71] and Idamycin [72] have also shown potential antitumor activity in HNSCC cell lines and xenograft models. Furthermore, Fedratinib has also been reviewed by Geiger *et al.* for HNSCC treatment that targets the janus kinase pathway involving in the activation of transcriptional factors and up regulation of cellular division and proliferation [73]. However, no studies were found that evaluated the efficiency of the three antineoplastic agents; Tivozanib, Mitoxantrone, and Selumetinib either using HNSCC or OSCC cell lines or model ogranisms.

The MoA of the well-characterized antineoplastic agents revealed inhibition of kinase proteins and signaling pathways involving in cell cycle and DNA replication. These candidates were indicated for a diverse range of cancers such as breast cancer, prostate cancer, acute lymphoid leukemia and renal cancer etc. The network analysis of antineoplastic agents and targeting genes resulted into 14 hub genes that were found enriched in the cellular proliferation, cell cycle and signaling. The enrichment of these pathways has been reported in several studies to involve in the tumorigenesis and progression of OSCC [55–57]. Five of the 14 hub genes; CCNB2, PLK1, CDK1, AURKA and CDC6 have been already reported as oncogenes [74]. In addition, the over expression of hub genes in the network were validated from Gepia2 online server by using HNSCC dataset. Hence, the genes perturbed by the well-characterized 12 antineoplastic agents showed significant inverse correlation with the DEGs in OSCC and could inhibit OSCC progression.

## Conclusion

Drug repositioning can be an efficient way for identification of candidate drugs or antineoplastic agents against OSCC. In this study, we used a systemic approach for identification of significant DEGs from multiple gene expression datasets of OSCC that regulated the biological pathways involving in the cancer progression. The correlation of the significant DEGs in OSCC and drug perturbation information from LINCS data resulted in 12 antineoplastic agents namely Teniposide, Palbociclib, Etoposide, Fedratinib, Tivozanib, Afatinib, Vemurafenib, Mitoxantrone, Idamycin, Canertinib, Dovitinib and Selumetinib. These antineoplastic agents are either FDA approved or are in advance clinical trial phases and have indications in various cancer types. The antineoplastic agents showed inhibitory interactions with the cluster of genes that were driving the phenotype of the OSCC. However, the efficiency of these antineoplastic agents in OSCC needs to be validated using *in vitro* and *in vivo* assays.

## Author Contributions

ASU and FFK contributed to the conception and design of the study; ASU undertook the analysis; and the drafting; FFK contributed to the critical review and editing of this manuscript.

## Acknowledgements

The authors declare no potential conflicts of interest with respect to the authorship and/or publication of this article. The authors are grateful to the Precision Medicine Lab grant by the National Centre for Big Data and Cloud Computing, Higher Education Commission and to CECOS University and Rehman Medical Institute, Peshawar, Pakistan for their generous support to the Lab.

## Conflict of interest

None declared.

## Notes

### Competing Interest Statement

The authors have declared no competing interest.

### Summary of Updates

Correct the author list.

